# Endorsement and Phylogenetic Analysis of some Fabaceae Plants based on DNA Barcoding gene *MatK*

**DOI:** 10.1101/2021.07.27.454001

**Authors:** Nader R. Abdelsalam, Mohamed E. Hasan, Samar M.A. Rabie, Houssam El-Din M.F. El-wakeel, Amera F. Zaitoun, Rehab Y. Ghareeb, Aly Z. Abdelsalam, Hesham M. Aly, Amira A. Ibrahim, Alaa A. Hemeida

## Abstract

DNA barcodes have been considered as a tool to facilitate species identification based on their simplicity and high-level accuracy compression to the complexity and subjective biases linked to morphological identification of taxa. MaturaseK gene “*MatK”* of the chloroplast is very crucial in the plant system which is involved in the group II intron splicing. The main objective of this current study is determining the relative utility of the “*MatK”* chloroplast gene for barcoding in fifteen legume trees by both single region and multiregional approaches. The chloroplast “*MatK”* gene sequences were submitted to GenBank and accession numbers (GenBank: LC602060, LC602154, LC602263, LC603347, LC603655, LC603845, LC603846, LC603847, LC604717, LC604718, LC605994, LC604799, LC605995, LC606468, LC606469) were obtained with sequence length ranging from 730 to 1545 nucleotides. These DNA sequences were aligned with database sequence using PROMALS server, Clustal Omega server and Bioedit program. Also, the maximum likelihood and neighbor-joining algorithms for phylogenetic reconstruction using the MEGA-X program were employed. Overall, these results indicated that the phylogenetic tree analysis and the evolutionary distances of an individual dataset of each species were agreed with a phylogenetic tree of all each other consisting of two clades, the first clade comprising *(Enterolobium contortisiliquum, Albizia lebbek), Acacia saligna, Leucaena leucocephala, Dichrostachys Cinerea, (Delonix regia, Parkinsonia aculeata), (Senna surattensis, Cassia fistula, Cassia javanica)* and *Schotia brachypetala* were more closely to each other, respectively. The remaining four species of *Erythrina humeana, (Sophora secundiflora, Dalbergia Sissoo, Tipuana Tipu)* constituted the second clade. Therefore, *MatK* gene is considered promising a candidate for DNA barcoding in plant family Fabaceae and providing a clear relationship between the families. Moreover, their sequences could be successfully utilized in single nucleotide polymorphism (SNP) or part of the sequence as DNA fragment analysis utilizing polymerase chain reaction (PCR) in plant systematic.

## 1. Introduction

Fabaceae is considering a large and economically vital family of flowering plants which is usually known as the legume family [1-3]. The Fabaceae family, which includes 730 genera more than 19,400 species, is the second-largest family of medicinal plants and the third largest family of flowering plants, with over 490 medicinal plant species [4-6]. Documentation of the Mediterranean legume crops depending on morphological characteristics has shown tricky and much impossible [7-9]. Using a DNA-based technique would offer accurate knowledge and facilitate the discrimination of the species. DNA barcoding is new, efficient, quick, low-cost, and standard technique for the fast identification and evaluation of plant and animal species based on DNA sequence from a small fragment of the whole genome in a rapid, accurate [10-14]. Also, DNA barcoding can help to detect species, quick identification of any species that are possibly novel to science and to report the essential ecological and evolutionary questions as a biological instrument [15-21]. DNA barcodes are frequently promoted for their facility to enhance public and non-specialist access to scientific information and new knowledge [22]. Short DNA sequences in DNA barcoding are used to identify the diversity between plant and animal species as molecular markers [23], also, it’s used in an assignment of the unknown samples to a taxonomic group, and in-plant biodiversity documentation [24]. DNA barcoding is a potential tool to detect an error in identifying species because similarity-based approaches using DNA barcoding combined with morphology would solve the misidentification based on morphology [25, 26]. DNA barcoding could help decrease the limitations of morphological characteristics and hurry up plant and animal species identification since it can detect the organisms at any stage of growth. DNA barcodes are intended to generate a distributed community resource of DNA sequences that would be used to identify or classify species by taxonomic categorization [27]. The usage of DNA barcodes as a tool for plant/ animal identification is based on the establishment of high-value reference databases of sequence [28] which cannot always distinguish between closely related species of land plants or fungi.

The matK gene (1500 bp in length), located inside the intron of the trnK and codes for protein maturase, which is involved in Group-II intron splicing. This gene has a high substitution rate [3], a huge fraction of variance in nucleic acid levels between the first and second codon positions, a low transition and or/transversion ratio, and mutationally conserved areas. Previous data were utilized to identify the molecular markers, which were used to identify the genus/species of these taxa, to provide valuable information for both conventional and molecular studies [8]. During this current study, we evaluated the capacity and the efficiency of the normal plant barcode marker *MatK* for documentation and identification of 45 plant specimens belonging to 15 species of Fabaceae plant species to study the useful annotation, homology modeling and sequence analysis to permit an additional efficient use of these sequences between different plant species.

## 2. Materials and methods

### 2.1. Plant materials

Forty-five samples (three replicates for each species), which belonged to 15 species found in Fig (1) were collected from Antoniadis Garden’s (N 29” 56’55, E 18” 12’31) between July 2019 and January 2020 and combined their sequences with those available from GenBank.

### 2.2. Extraction and sequencing of DNA from samples

Total genomic DNA was extracted from fresh leaves tissue using i-genomic plant DNA extraction Mini kit @ iNtron biotechnology according to the Plant Genomic DNA Kit procedure (iNtRON Bio Co., South Korea). PCR of the *MatK* regions were conducted out in Techne Flexigene PCR Thermal Cycler programmed for 30 cycles as follows: 94°C/5 minutes (1 cycle); 94°C/45 sec, 50°C/45 sec, 72°C/45 sec (30 cycles); 72°C/7 minutes (1 cycle); 4°C (infinitive). The designed common primers and reaction conditions of the *MatK* region is F: 5’-CGTACAGTACTTTTGTGTTTACGAG-3’ (Tm, 53.9 & GC%, 40), R: 5’-ACCCAGTCCATCTGGAAATCTTGGTTC-3’ (Tm, 60.4 & GC%, 48). The PCR products were electrophoresed on a 1.0% agarose gel with 1X TAE buffer containing 0.5 g/mL ethidium bromide. PCR products were purified with Mini kit @ iNtron Biotechnology Purification kits before being sequenced with a DNA sequencer (Applied Biosystems® 3500 and 3500xL Genetic Analyzers) and a BigDye Terminator version 3.1 Cycle Sequencing RR-100 Kit using the dideoxynucleotide chain termination method (Applied Biosystems). The sequences were submitted to DDBJ/EMBL/GenBank database. Generic and species data was achieved from the taxonomy database of the National Centre for Biotechnology Information (NCBI).

### 2.3 Sequence analysis

The sequencing results analysis has been done for a unique grouped dataset that contains all of the Fabaceae plant species for which sequences are available in GenBank to identify inter-species and inter-generic variation. *Fabaceae* species sequences of *MatK* were retrieved from NCBI in Fasta format. Multiple sequence alignments of the *MatK* gene were conducted from different species applying the PROMALS server [29], Clustal Omega server [30], the BIOEDIT software [31] and MEGA-x [32] which are offline software that conducts optimal sequence alignment to find the conserved area. Comparing to the greatest alignment methods with development for distantly related sequences the “PROMALS” is up to 30% more accurate. Clustal Omega server is a new multiple sequence alignment software that generates alignments between three or more sequences using seeded guide trees and HMM profile-profile methods. The “BIOEDIT” software is a user-friendly biological sequences alignment editor that aims to provide fundamental functions for editing, aligning, manipulating, and analyzing protein sequences.

#### 2.3.1 Molecular evolution and phylogenetic analysis

The Neighbor Joining method was used to deduce the evolutionary narrative. Finding the topology and branch length of the tree that will offer the best chance of detecting amino acid sequence in current data is the approach for constructing the phylogenetic tree using maximum likelihood. So, for phylogenetic evaluation Mafft server [33], Clustal Omega server and “MEGA-x” software were applied. MEGA was used to analyze the sequencing data using the neighbor-joining technique and Unweighted Pair Group Mean Average “UPGMA.” The “DNADIST” software of “PHYLIP” was used to calculate distances. NJ plot was used to do bootstrapping and decay analysis. MEGA determined parsimony analyses and different clades.

## 3. Results

### 3.1. DNA extraction and PCR amplification

The quality of the obtained DNA was detected 1% agarose gel electrophoresis as seen in Fig 2. The results indicated that there is no fragmentation was observed in extracted DNA. The quantity of extracted DNA samples was determined by using Nanodrop Spectrophotometer and the concentration ranged from 30 -50 ng/μl. The extracted DNA was directly used in PCR amplification as found in Fig 3 for the *MatK* gene (900 bp). Development in DNA sequencing methods has allowed us to describe the genomes of numerous organisms quickly. Evaluations of the DNA sequences of several species are providing useful knowledge about their taxonomy, gene makeup, and utilization. In the current study using DNA sequence polymorphisms of the chloroplast, *MatK* gene is much more variable than many other genes. From Fifteen plant species belong to different genera of the same family (Fabaceae), we organized a study to contribute to the knowledge of the major evolutionary relationship between the studied plant genus and species (clades) and discussed the application of *MatK* for molecular evolution. The chloroplast *mat*K marker was more useful as DNA markers.

**Fig (1).**
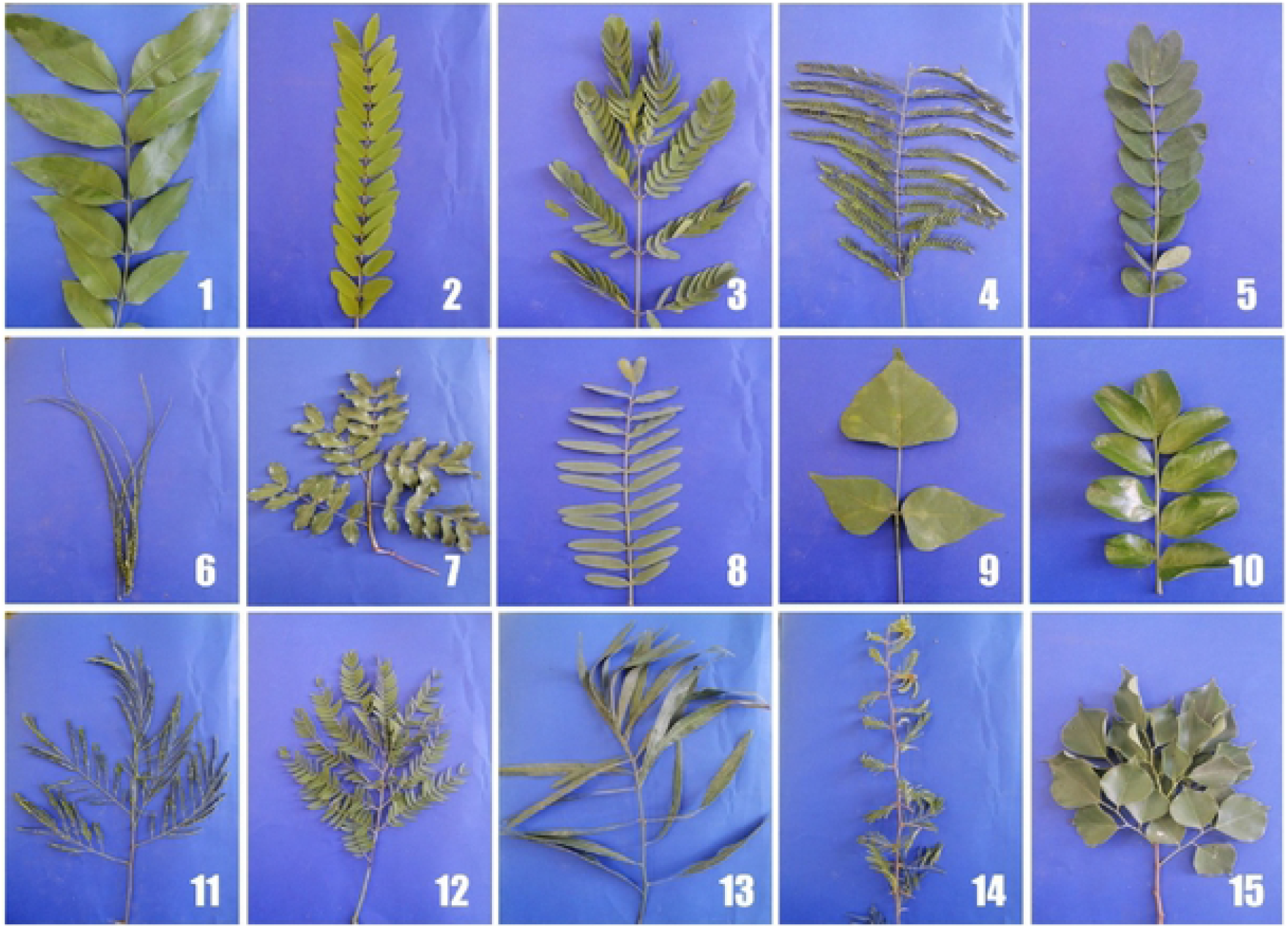
showing different leaves in fifteen Fabaceae plants (1)*Cassia fistula*, (2) *Cassia javinca*, (3) *Albizia lebbek*, (4) *Delonix regia*, (5) *Senna surattensis*, (6) *Parkinsonia aculeata*, (7) *Schotia brachypetala*, (8) *Tipuana tipu*, (9) *Erythrina humeana*, (10) *Sophora secundiflora*, (11) *Leucaena leucocephala*, (12) *Enterolobium contortisiliquum*, (13) *Dichrostachys cinerea*, (14) *Acacia saligna and* (15) *Dalbergia sissoo*

**Fig (2).**
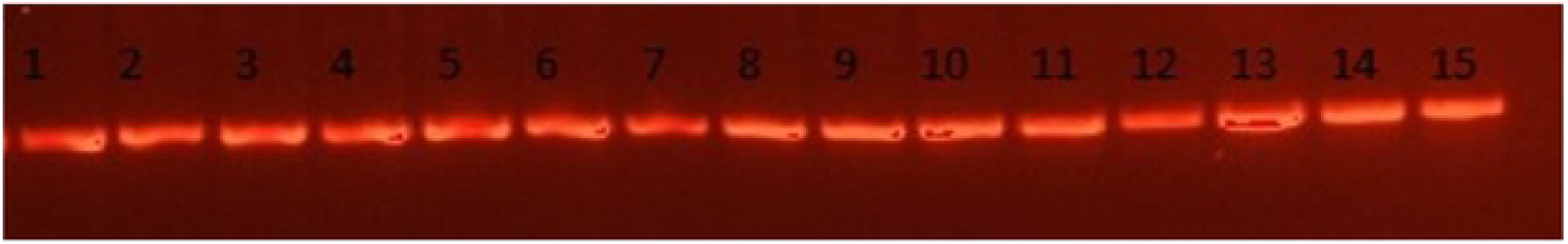
Agarose gel electrophoresis for extracted DNA from samples (1) *Cassia fistula*, (2) *Cassia javinca*, (3) *Albizia lebbek*, (4) *Delonix regia*, (5) *Senna surattensis*, (6) *Parkinsonia aculeata*, (7) *Schotia brachypetala*, (8) *Tipuana tipu*, (9) *Erythrina humeana*, (10) *Sophora secundiflora*, (11) *Leucaena leucocephala*, (12) *Enterolobium contortisiliquum*, (13) *Dichrostachys cinerea*, (14) *Acacia saligna and* (15) *Dalbergia sissoo*

**Fig (3).**
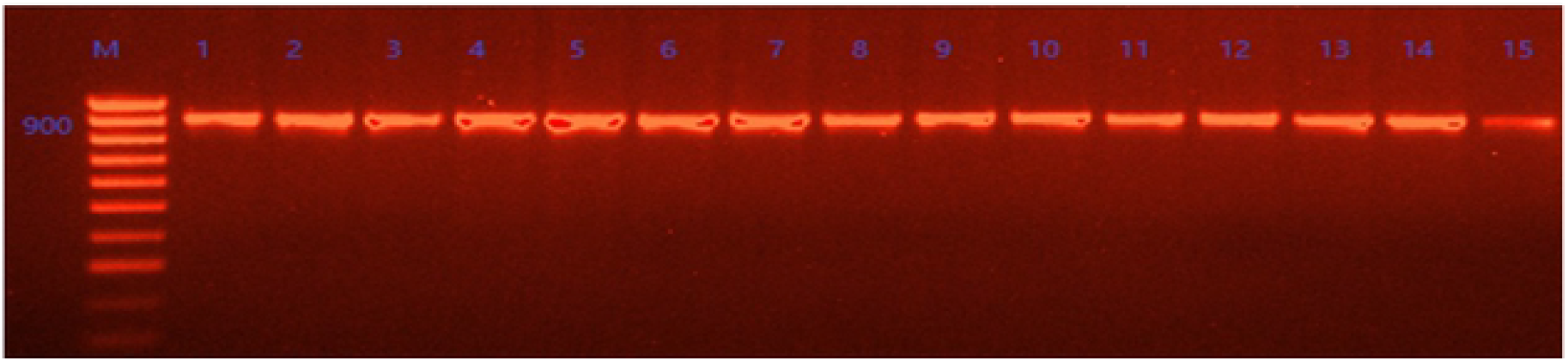
Agarose gel electrophoresis for amplified samples by using the primer *MatK* for the samples (1) *Cassia fistula*, (2) *Cassia javinca*, (3) *Albizia lebbek*, (4) *Delonix regia*, (5) *Senna surattensis*, (6) *Parkinsonia aculeata*, (7) *Schotia brachypetala*, (8) *Tipuana tipu*, (9) *Erythrina humeana*, (10) *Sophora secundijlora*, (11) *Leucaena leucocephala*, (12) *Enterolobium contortisiliquum*, (13) *Dichrostachys cinerea*, (14)*Acacia saligna and* (15) *Dalbergia sissoo*

**Table 1.**
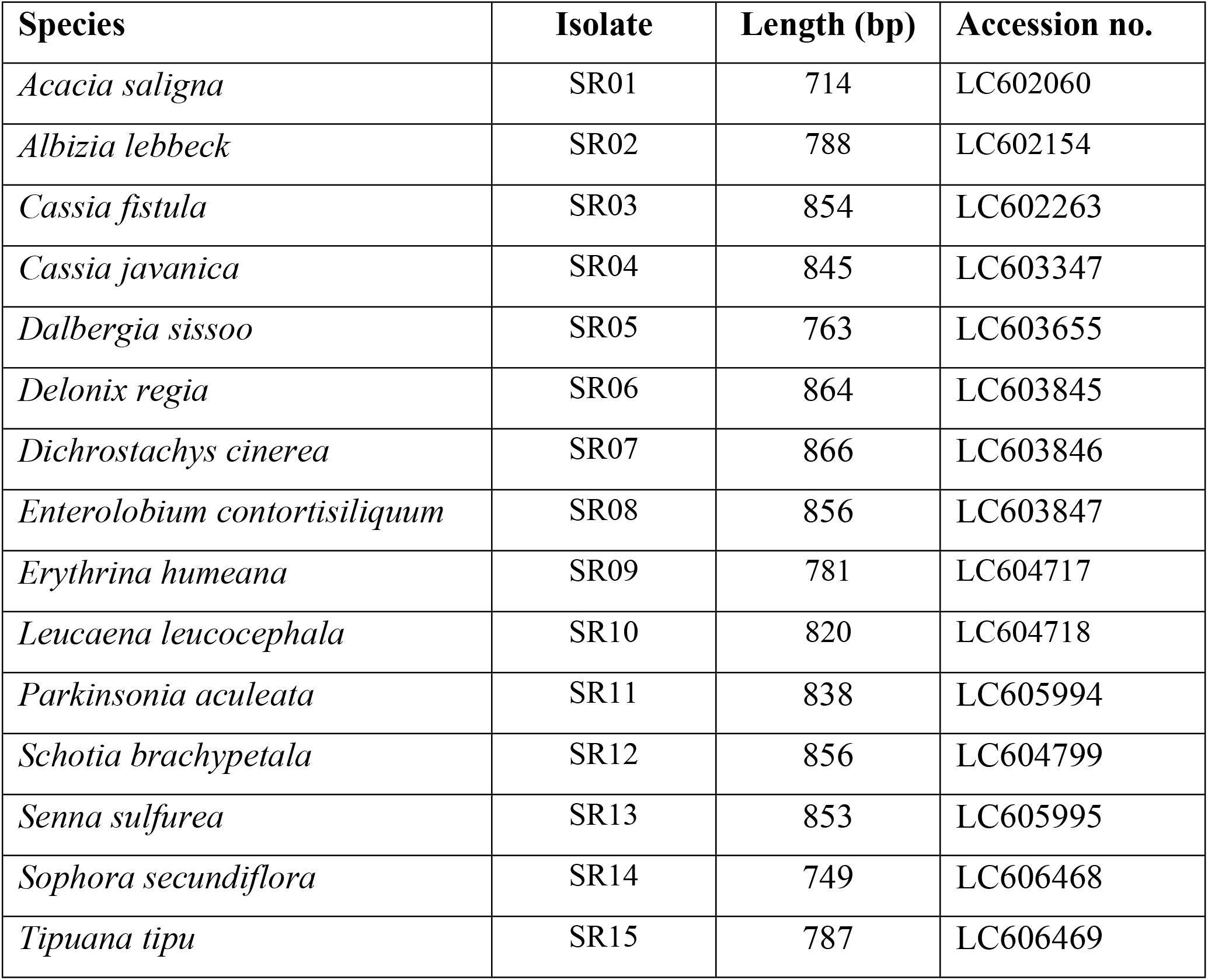
Fifteen taxa, their isolates, the sequence length (bp), and assigned Accession numbers of each plant used in this study.

| Species | Isolate | Length (bp) | Accession no. |
| --- | --- | --- | --- |
| *Acacia saligna* | SR01 | 714 | LC602060 |
| *Albizia lebbeck* | SR02 | 788 | LC602154 |
| *Cassia fistula* | SR03 | 854 | LC602263 |
| *Cassia javanica* | SR04 | 845 | LC603347 |
| *Dalbergia sissoo* | SR05 | 763 | LC603655 |
| *Delonix regia* | SR06 | 864 | LC603845 |
| *Dichrostachys cinerea* | SR07 | 866 | LC603846 |
| *Enterolobium contortisiliquum* | SR08 | 856 | LC603847 |
| *Erythrina humeana* | SR09 | 781 | LC604717 |
| *Leucaena leucocephala* | SR10 | 820 | LC604718 |
| *Parkinsonia aculeata* | SR11 | 838 | LC605994 |
| *Schotia brachypetala* | SR12 | 856 | LC604799 |
| *Senna sulfurea* | SR13 | 853 | LC605995 |
| *Sophora secundiflora* | SR14 | 749 | LC606468 |
| *Tipuana tipu* | SR15 | 787 | LC606469 |

### 3.2. Phylogenetic analysis of collected plants

Numerous sequence alignments showed that there are varying numbers of “Indels” in the gene *MatK*. Using the neighbor-joining method, UPGMA and maximum likelihood, the evolutionary distances for the 15 plant species were recognized into individual clades. The alignment of *MatK* gene of *Acacia saligna* nucleotide sequences showed 769 variable sites and 571 parsimony sites, the overall mean distance is 2.85. The combined tree showed two groups or cladograms and they represented as follows: Group I include *Acacia saligna* was closely related to different species belong to other genera of the same family (Fabaceae) such as *Enterolobium, Pararchidendron, Archidendron, Samanea, Hydrochorea, Balizia* and *Abarema* (Fig 4). Also, *Acacia* comprising other species were closely arranged but distinguished to different genera such as *Falcataria, Pararchidendron* and *Lacacia*. In addition, the aligned *MatK* dataset was 793 nucleotide sites long, of which 102 sites were potentially parsimony informative. Consequently, *Enterolobium contortisiliquum* is more closely related to different species of genus *Acacia* according to phylogenetic analysis using maximum likelihood (Fig 5). The length of *MatK* varies from 750 bp in *Albizia lebbek* (the smaller length of *MatK* gene for these species is due to incomplete sequencing, which was retrieved from GenBank) to 813 in different genera *(Enterolobium, Acacia, Senegalia, Cojoba, Samanea, Hydrochorea, Balizia* and *Abarema)*. Maximum likelihood and Neighbor-joining analysis of the dataset resulted in a tree with two groups. The clades established in the trees were mainly mixtures of numerous species. Consequently, creating a local barcode database will be useful for a broad range of potential ecological purposes, involving the building of community phylogenies [34]. GroupI has three clusters comprising several genera (*Albizia, Enterolobium, Mariosousa, Archidendron, Samanea, Balizia*, and *Abarema)*. Otherwise, group II has one genera acacia which is the most closely related to our plant *Albizia lebbeck* according to *MatK* gene partial cds (Fig 6). The arrangement of *MatK* gene of *Acacia saligna* nucleotide sequence revealed 649 varying sites and 359 parsimony sites, the overall mean distance is 2.37 and the estimated Transition/Transversion bias (R) is 0.52.

**Fig 4:**
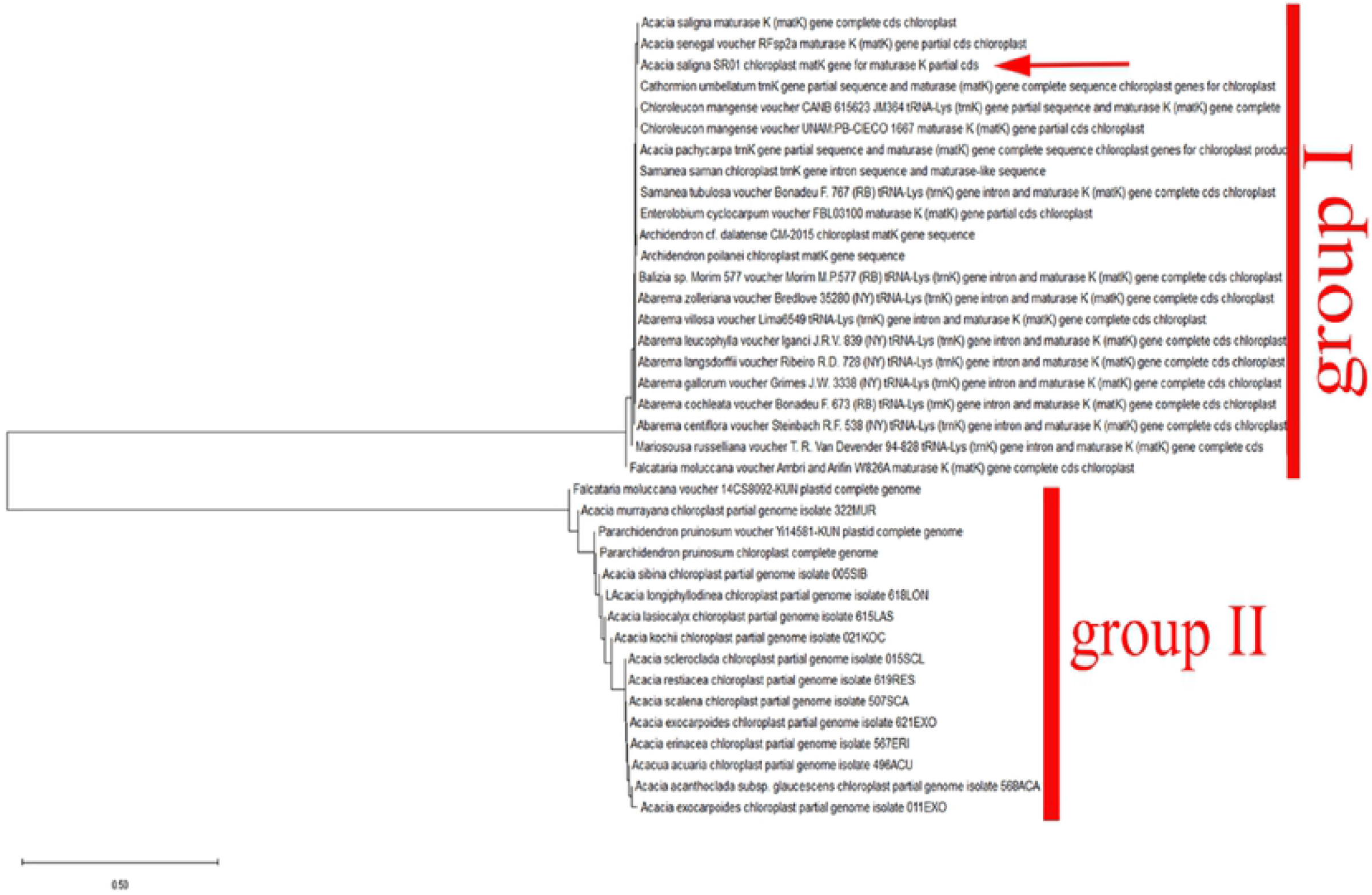
Phylogenetic tree analysis and the evolutionary distances of *Acacia saligna* were computed using the Maximum Likelihood technique using nucleotide sequences of the *MatK gene*.

**Fig 5:**
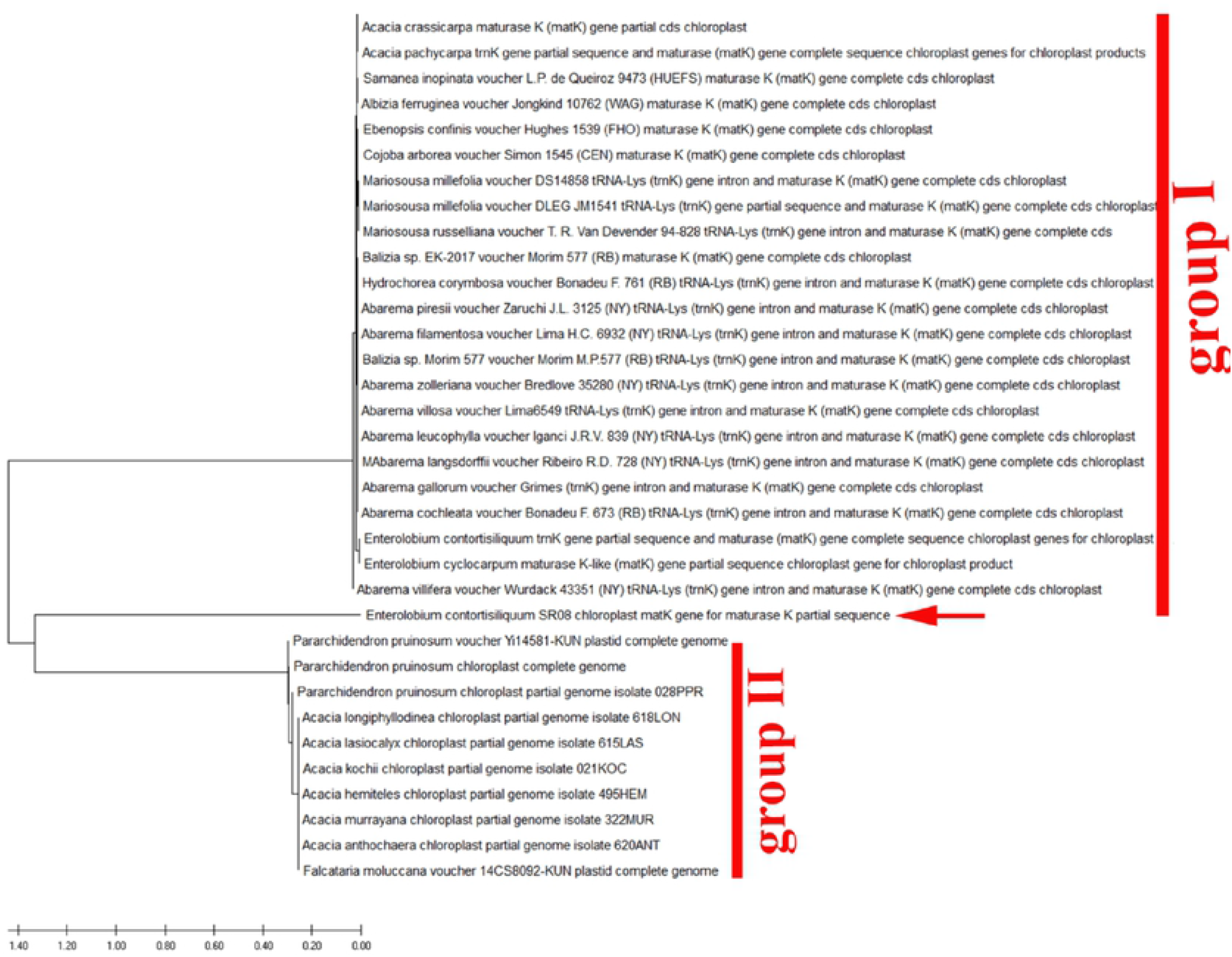
Phylogenetic tree analysis and the evolutionary distances of *Enterolobium contortisiliquum* were computed using the Maximum Likelihood technique using on nucleotide sequences of the *MatK gene*.

**Fig 6:**
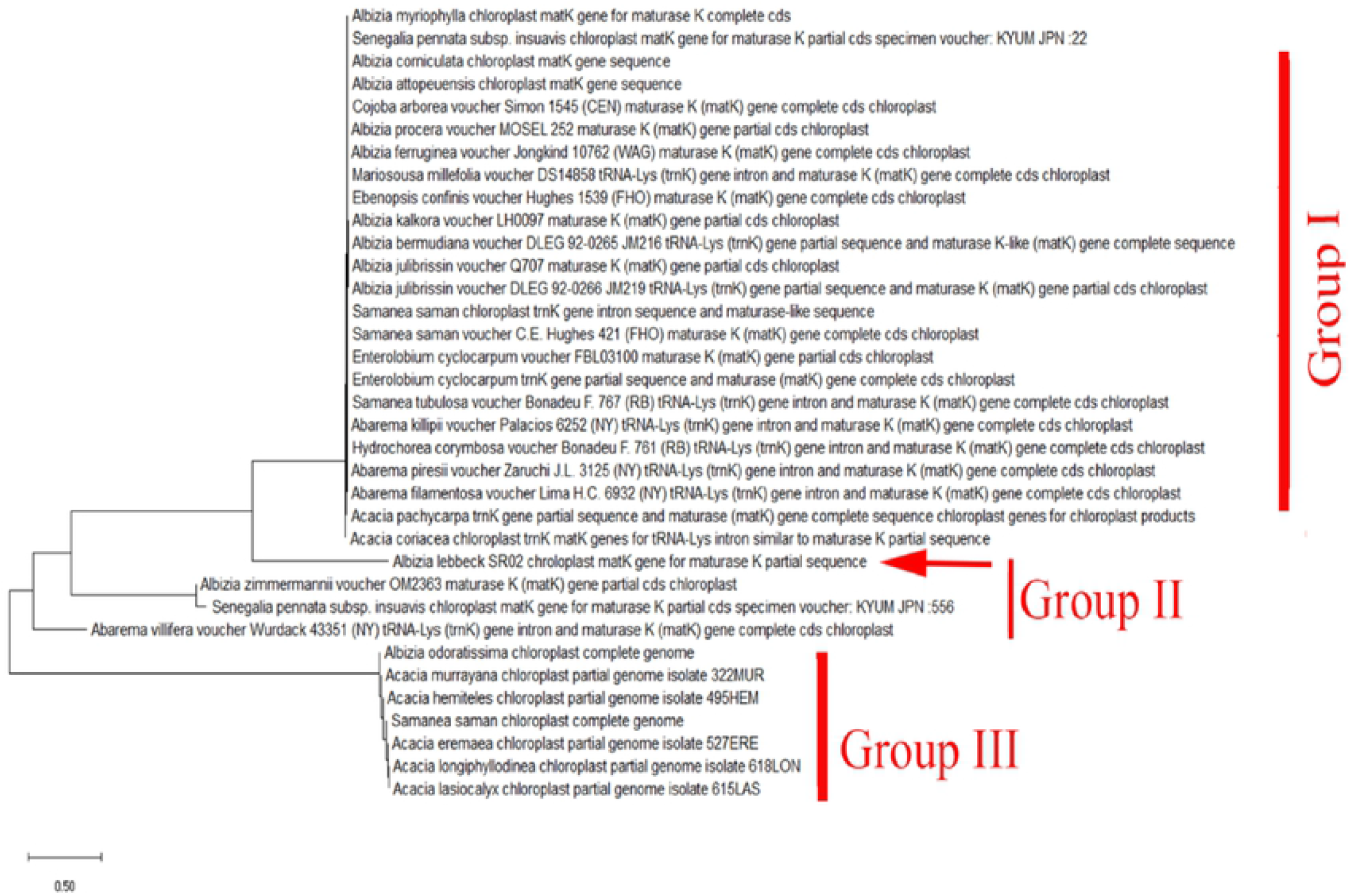
Phylogenetic tree analysis and the evolutionary distances of *Albizia lebbeck* were computed using the maximum likelihood technique using nucleotide sequences of the *MatK gene*.

Furthermore, depending on the phylogenetic analysis, the two genera *Cassia* and *Senna* with different species are closely related and more highly similar than any other studies species (Figs 7-9). The phylogeny tree was created using the neighbor-joining approach and the evolutionary distances were calculated employing the maximum composite likelihood approach. The combined trees showed that there are two groups, and they are as follows: Group I consisted of five clades representing different genera with different species such as (*Chamaecrista, Senna, Erytherophleum, Arapatiell* and *Dinizia*). Group II showed two branches: each one with many sub-branches containing five clades with different species of genus *Senna*. According to *MatK* gene sequence, the collected plants (*Cassia fistula* and Cassia *javanica*) revealed a high percentage of identity with different 7 species of genus senna having the same clade (Fig 7-8). Also, they are closely related to other different species of *Erytherophleum, Arapatiell*. On another hand, the sequence of *MatK* gene of collected *Senna surattensis* species has a high degree of similarity with many species in different genera in Fabaceae (Fig 9), and consequently this species used as a template to estimate the similarity between different species in Fabaceae family.

**Fig 7:**
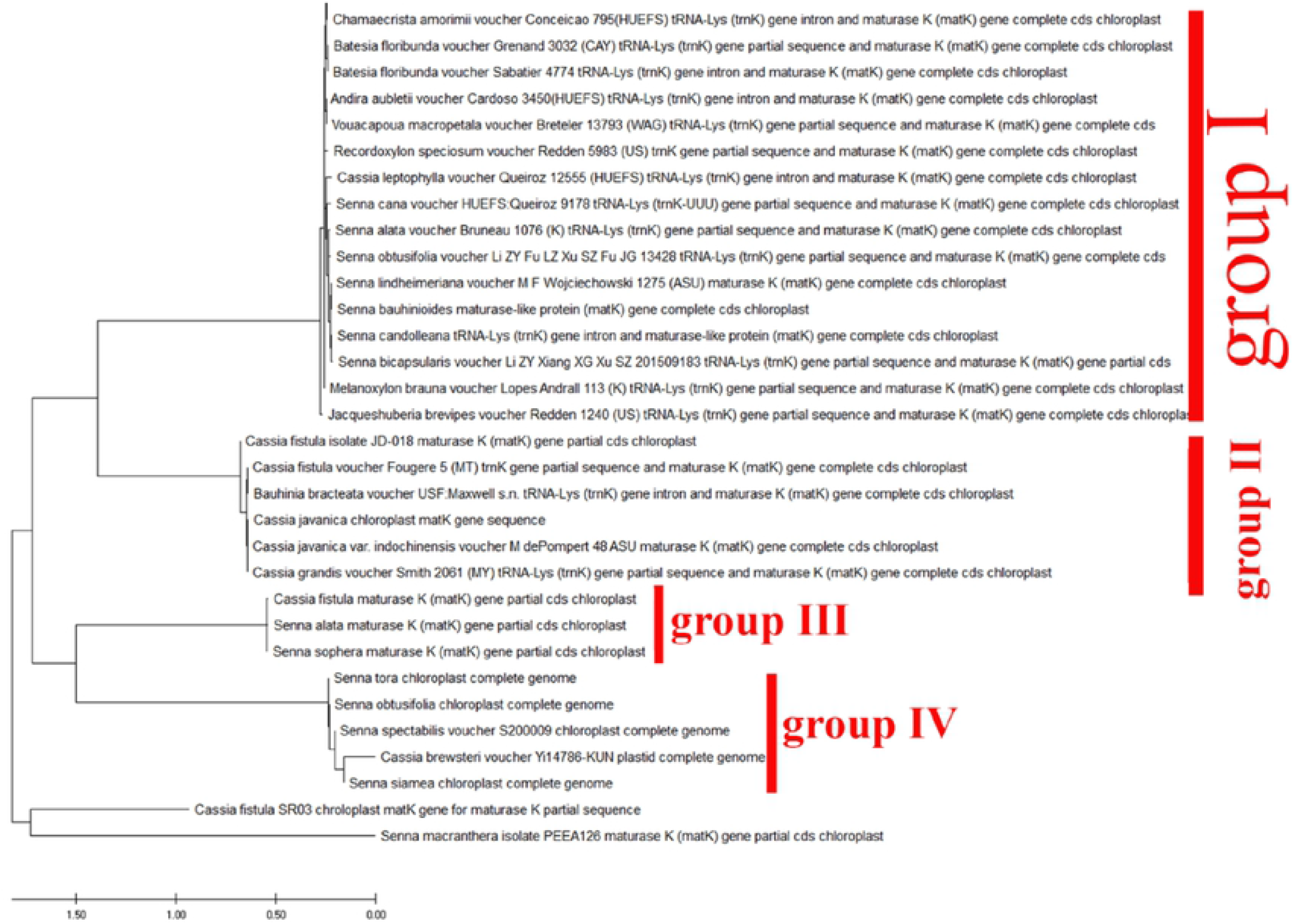
Phylogenetic tree analysis and the evolutionary distances of *Cassia fistula* using the Neighbor-Joining technique using nucleotide sequences of the *MatK* gene.

**Fig 8:**
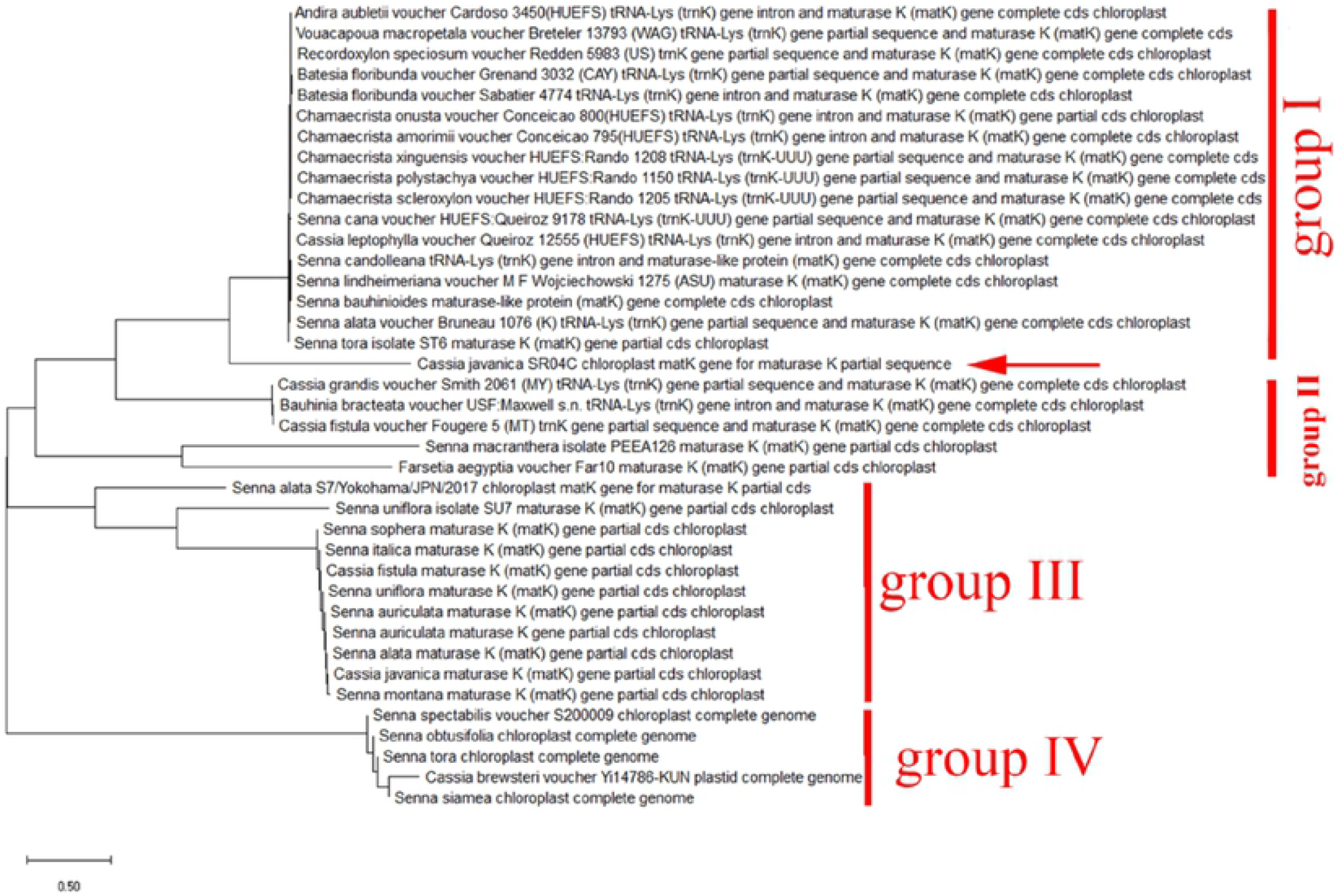
Phylogenetic tree analysis and the evolutionary distances of *Cassia javanica* using the Neighbor-Joining technique using on nucleotide sequences of the *MatK* gene.

**Fig 9:**
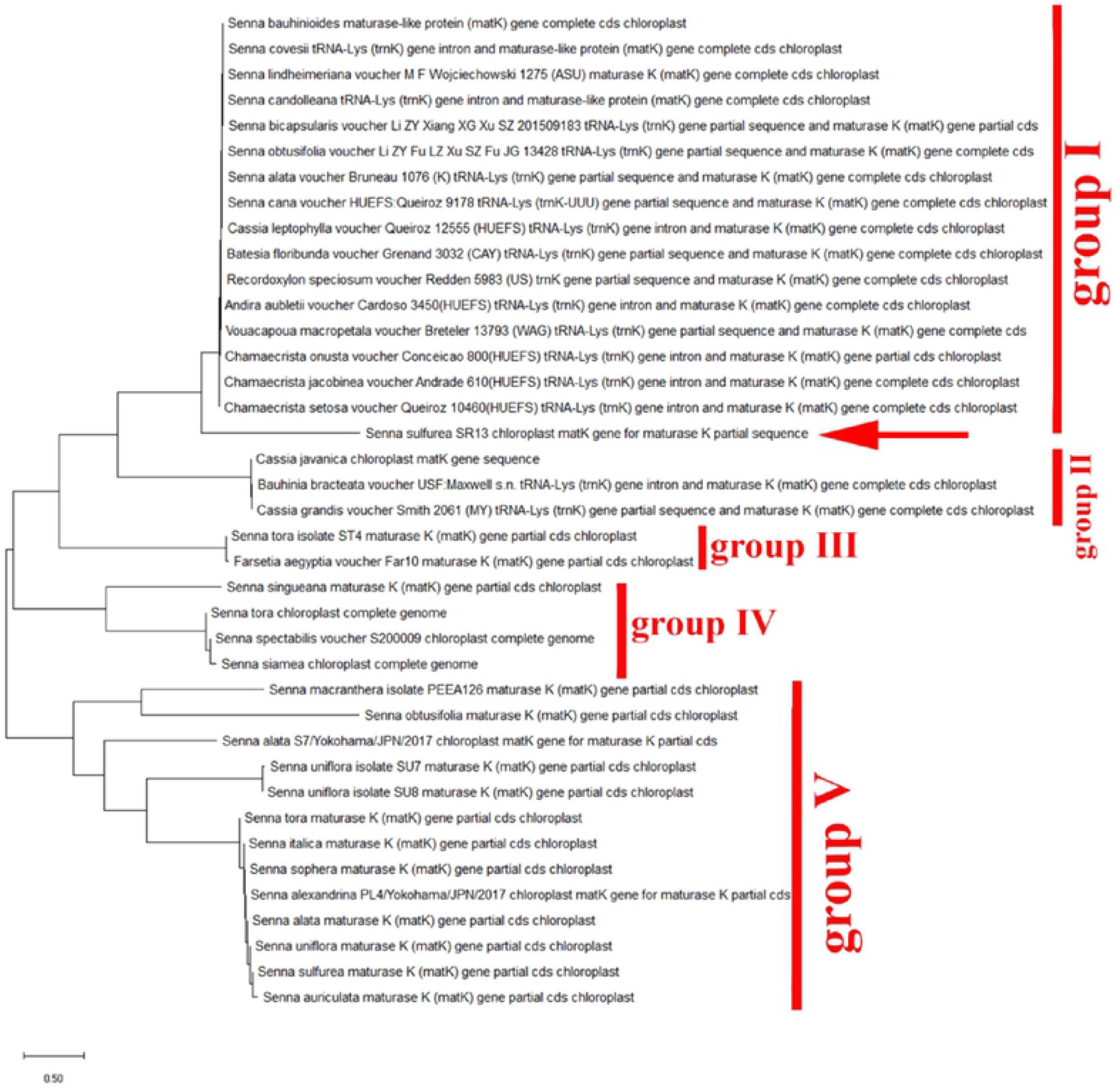
Phylogenetic tree analysis and the evolutionary distances of *Senna surattensis* using the

Results indicated the species of *Dalbergia sissoo, Erythrina humeana* and *Leucaena leucocephala* are more like other species of the same genus and less similar to species of other genera of the Fabaceae family (Fig 10 -12). Nevertheless, the *Delonix regia* is the more studied species having a good similarity to different species of different genera of the Fabaceae family. Polymorphism obtained from the DNA sequence indels or replacements of the *MatK gene* indicated that *Delonix regia, Umtiza listeriana, Dipty*chandra *aurantiaca, Moldenhawera blanchetiana, Schizolobium parahyba, Tachigali costaricensis, Arapatiella psilophylla* and *Parkinsonia Africana* were evolved from a Common ancestor (Fig 13). In addition, *Dichrostachys cinerea* is closely related to different species of genera *Leucaena, Senegalia, Falcataria and prosopis* (Fig 14). Furthermore, applying the same incremental method of informative sites starting at the 5^/^-end of the *MatK* gene, completely different results were found. The consensus tree of 15 most parsimonious trees demonstrated unresolved clades until 250 informative sites. At that point, 1 highly parsimonious tree was created, which was congruent with the topology of the stable tree achieved from the 3^/^-end. To recognize the greatest DNA barcode marker for species documentation and traceability, the value of genetic divergence for all the confirmed loci were calculated in each analyzed group at dissimilar taxonomic level and by considering only fresh morphologically identified samples.

**Fig 10:**
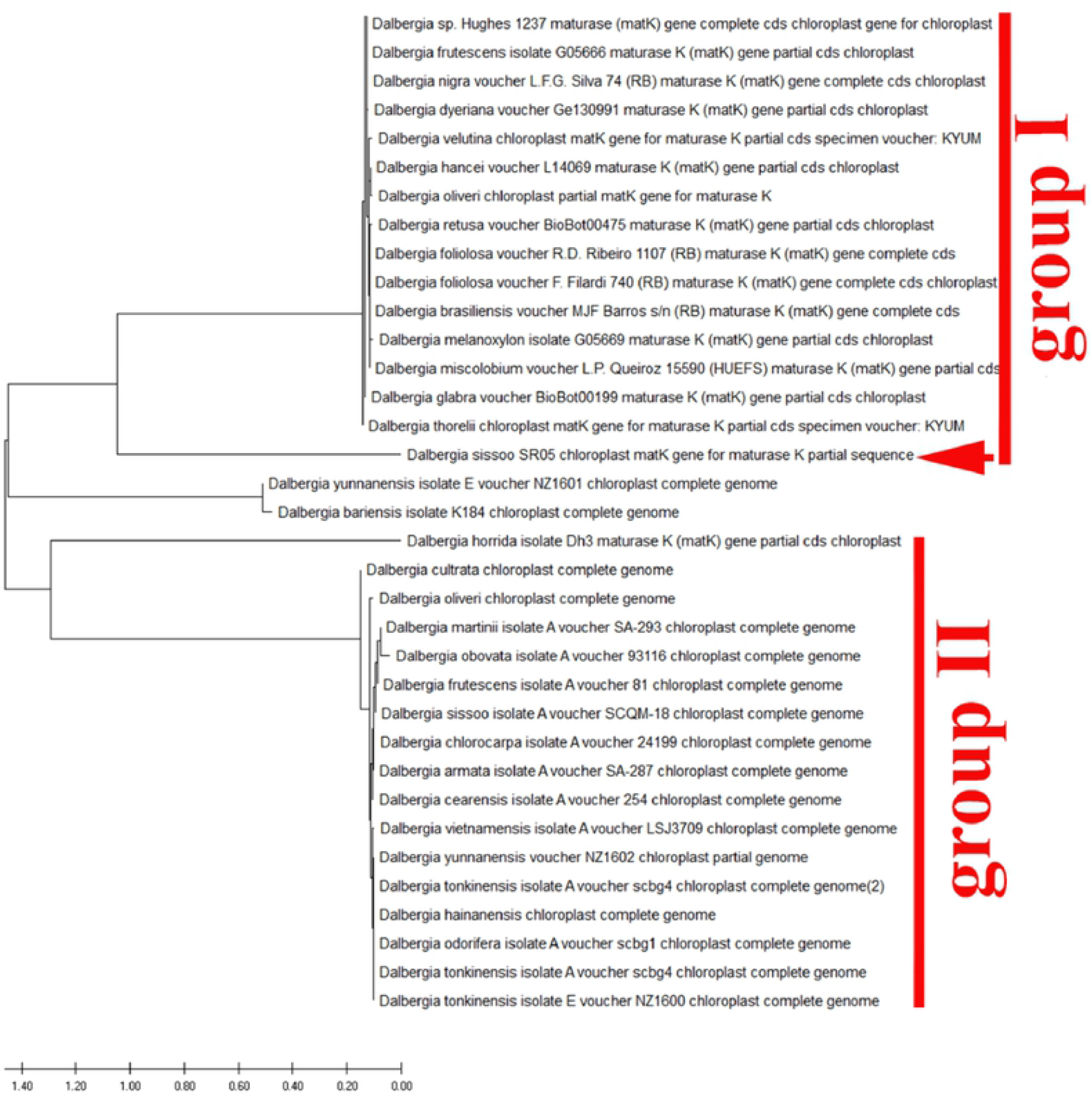
Phylogenetic tree analysis and the evolutionary distances of *Dalbergia Sissoo* using the Neighbor-Joining technique using nucleotide sequences of the *MatK* gene.

**Fig 11:**
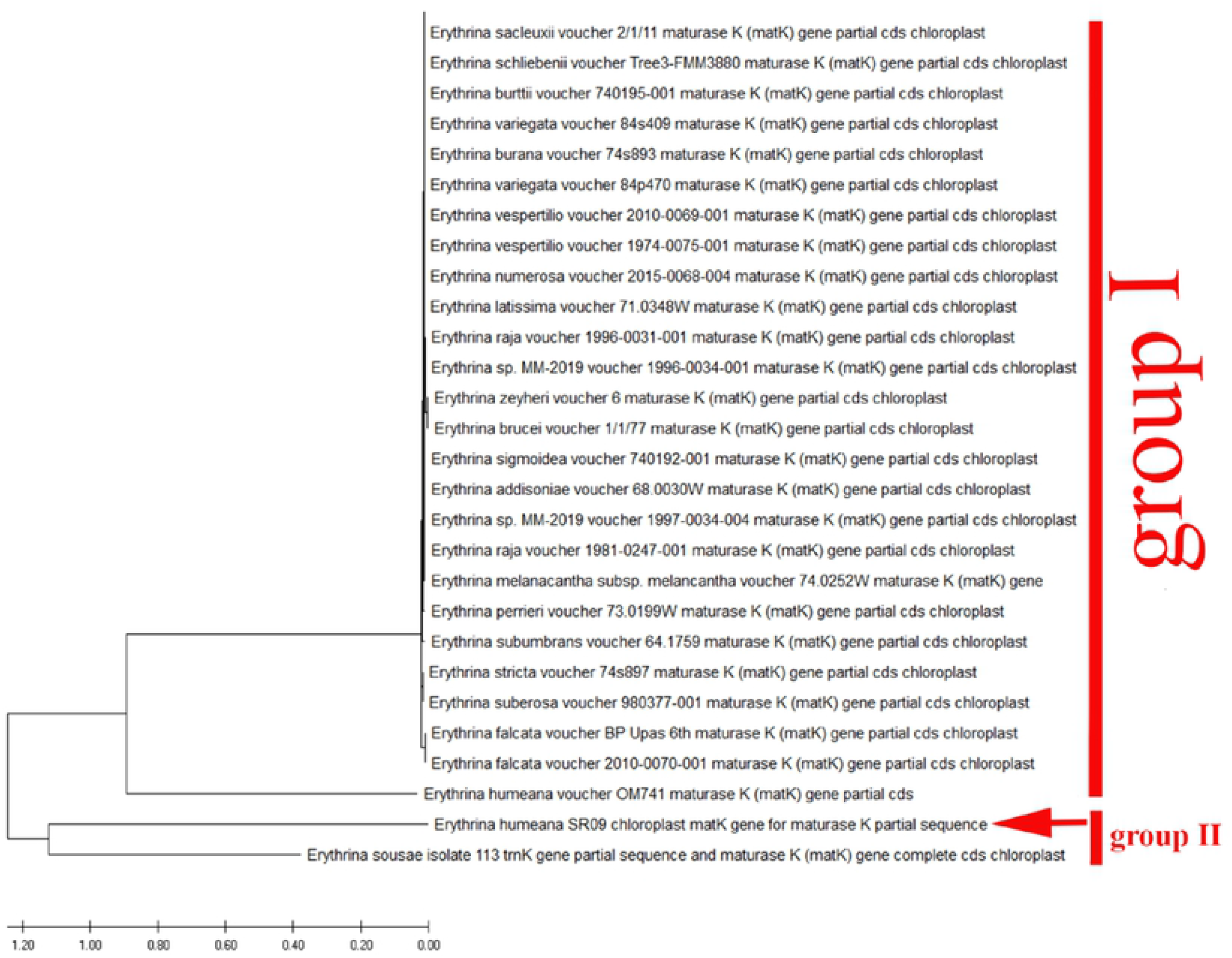
Phylogenetic tree analysis and the evolutionary distances of *Erythrina humeana* using the Maximum Likelihood technique using nucleotide sequences of the *MatK* gene

**Fig 12:**
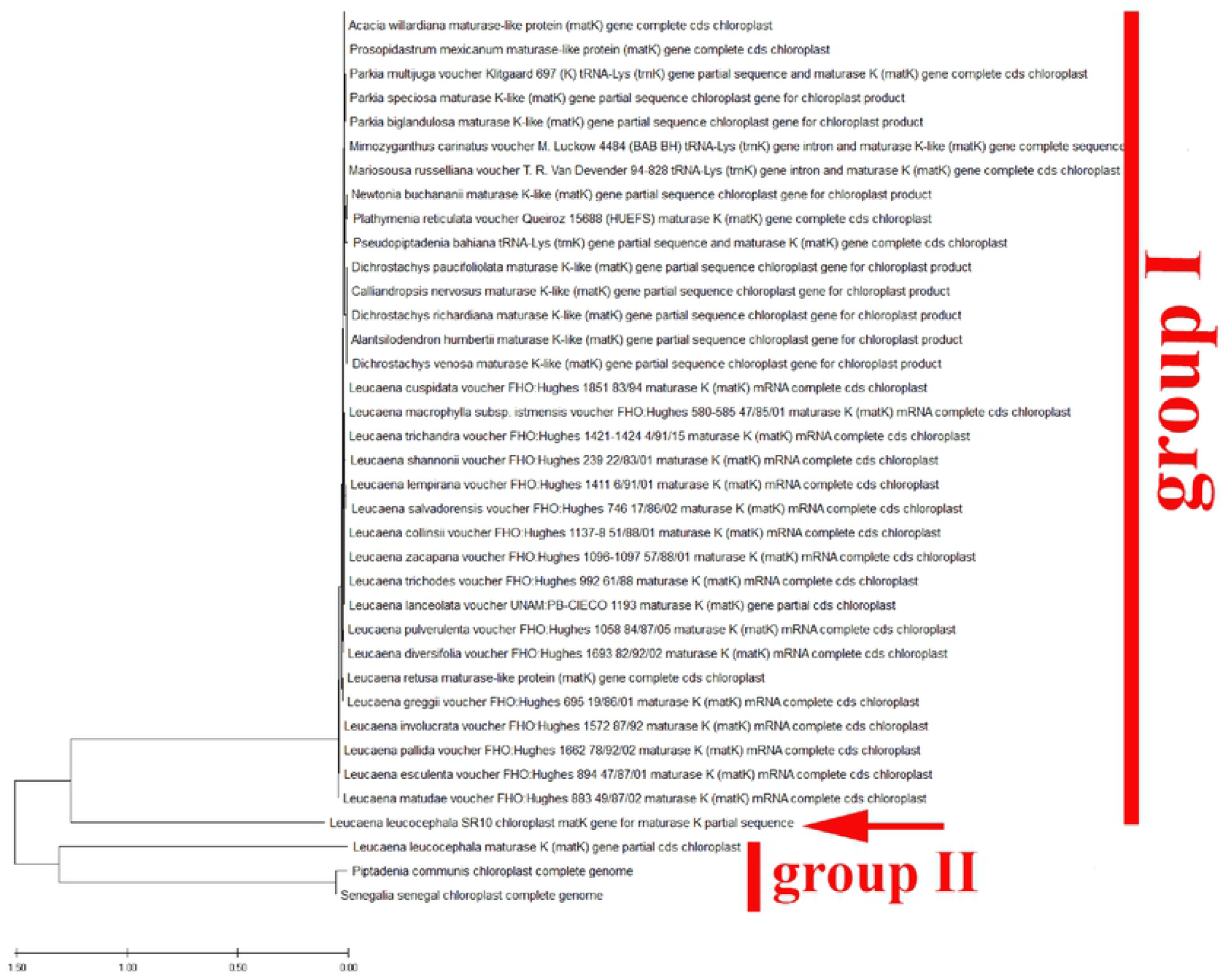
Phylogenetic tree analysis and the evolutionary distances of *Leucaena leucocephala* using the Neighbour Joining technique using nucleotide sequences of the *MatK* gene.

**Fig 13:**
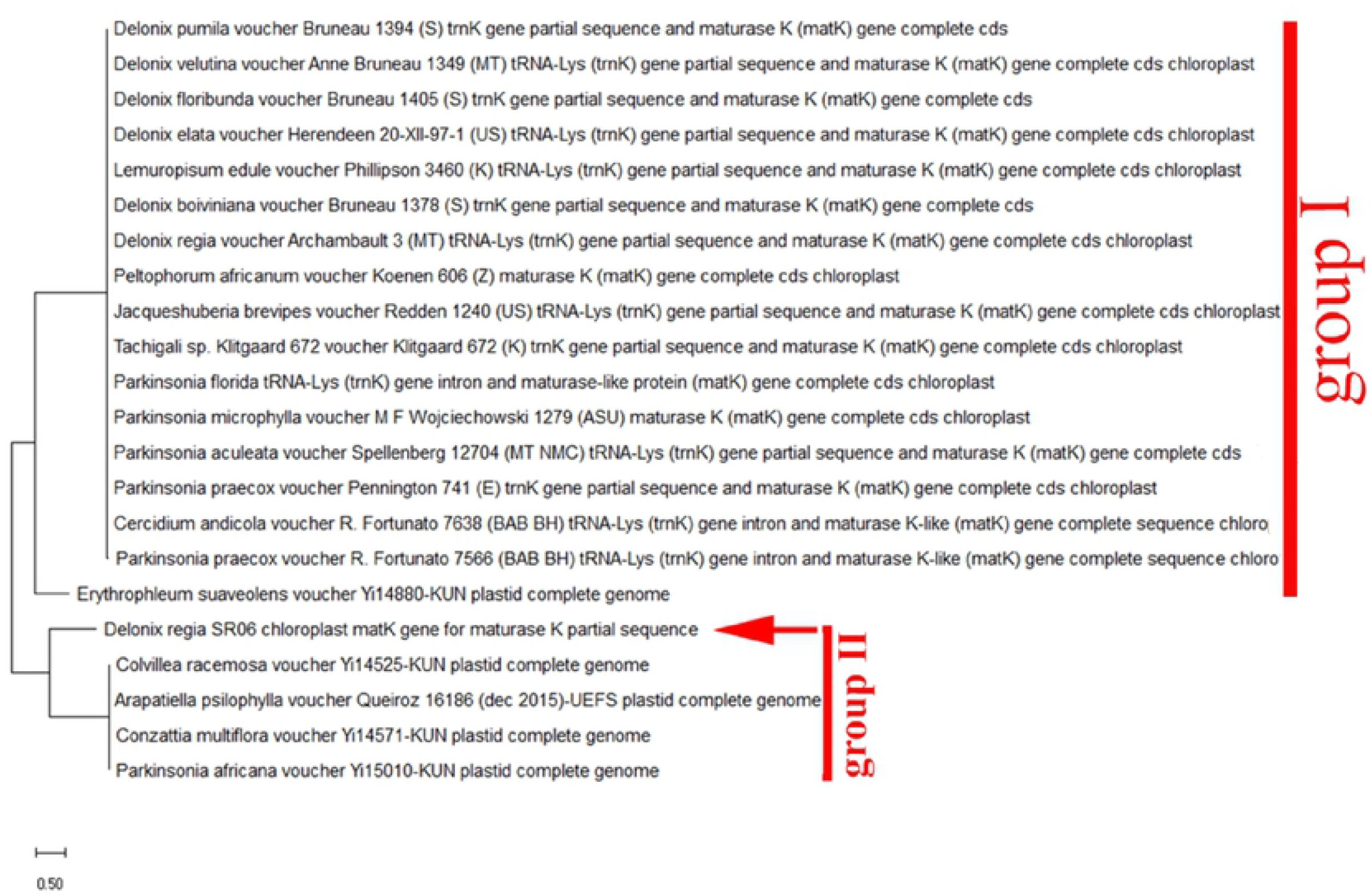
Phylogenetic tree analysis and the evolutionary distances of *Delonix*

**Fig 14:**
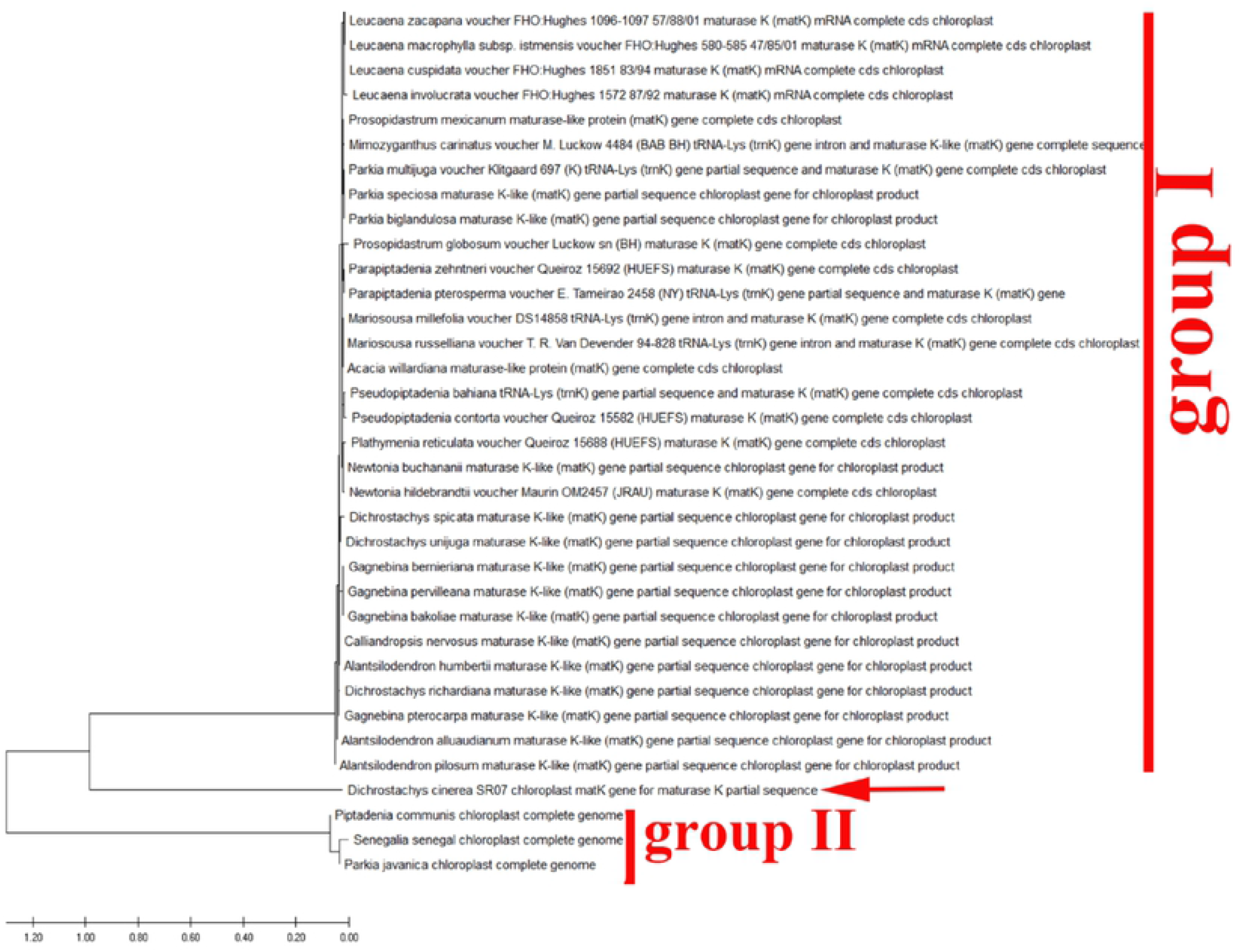
Phylogenetic tree analysis and the evolutionary distances of *Dichrostachys Cinerea*.

The current sequences showed little variations in the percentage of guanine plus cytosine content (% G+C) related to that in the sequences of *MatK*. In case of *MatK*, the nucleotide structure was biased toward the guanine and cytosine content (G+C) with frequencies were 30.4 to 34.8%, respectively. The NJ, ML, and MP analyses all resulted in comparable trees in each of the data sets. There are often variations between the trees from the various analyses involving non-resolution (polytomies). Analyses carried out on samples belonging to *Parkinsonia aculeata, Schotia brachypetala, Sophora secundiflora and Tipuana Tipu* indicated that the sequences divergences of marker *MatK* were clearly distinguished from other species of Fabaceae. Figs 15-18 showed phylogenetic clusters constructed using ML and NJ; The difference observed in *MatK* does separate several species; however, there is a wide range of intra-specific and inter-specific variation. The results of the sequence identification analysis reflected that the *Parkinsonia aculeata* was closely related and in the same clade with *Schizolobium parahyba, Diptychandra aurantiaca, Delonix regia, Conzattia multiflora and Colvillea racemose* (Fig 15). Furthermore, On the Neighbor-Joining Phylogram (Fig 16), the *Schotia* group is a sister taxon to the *Macrolobium* group and this observation is found in 50% of the most clade in this cladistic analysis (Fig 16).

**Fig 15:**
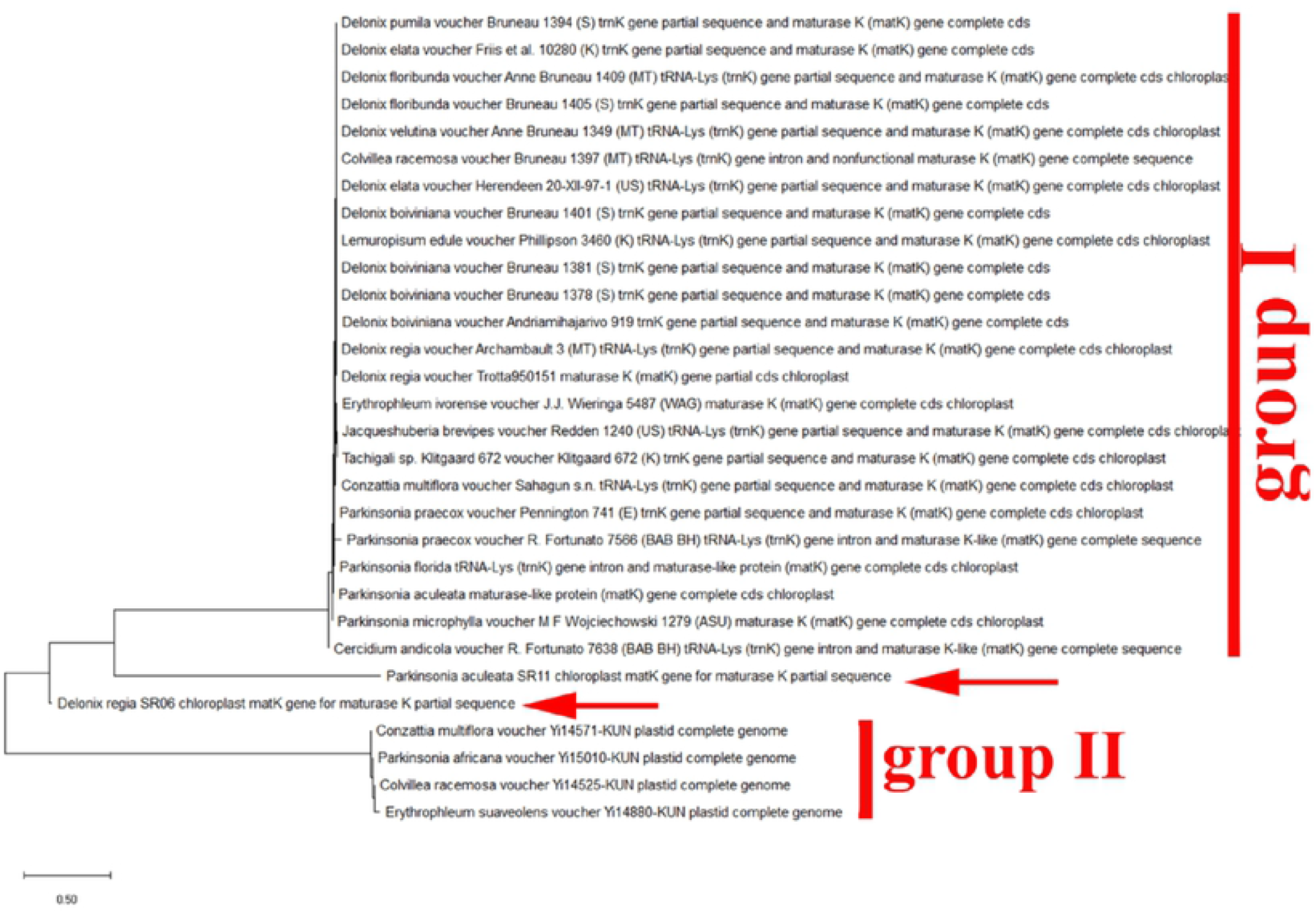
Phylogenetic tree analysis and the evolutionary distances of *Parkinsonia aculeata*

**Fig 16.**
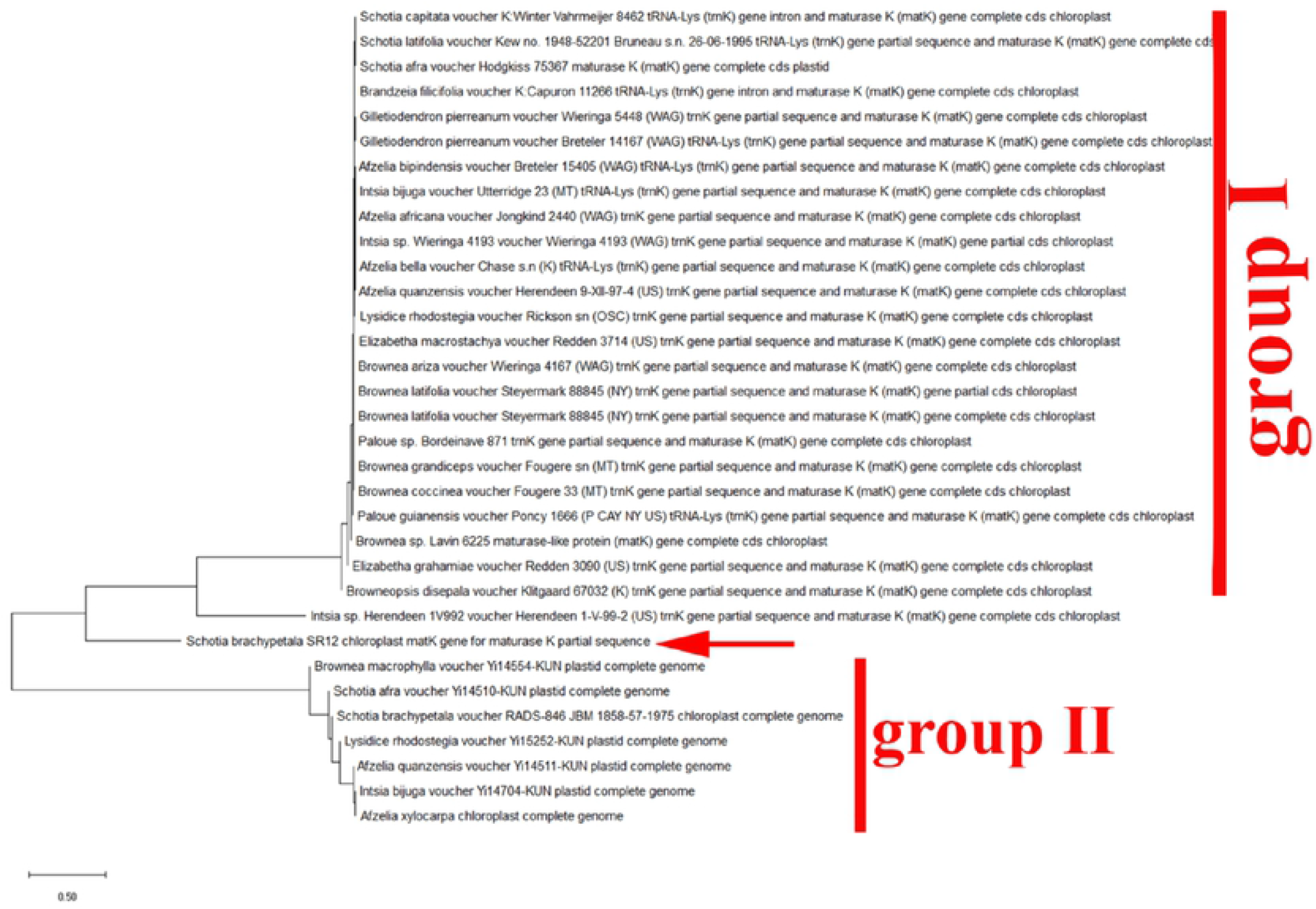
Phylogenetic tree analysis and the evolutionary distances of *Schotia brachypetala*

The last two members i.e., *Sophora secundiflora* and *Tipuana Tipu* produced an independent clade and confirmed the ambiguous position relative to the other genera of Fabaceae based on the combined cladistic analysis data from chloroplast DNA restriction sites and morphology. *Sophora secundiflora* shared a common ancestor with *Angylocalyx braunii, Zollernia splendens, Ormosia xylocarpa and Dermatophyllum secundiflorum* (Fig 17). Also, *Tipuana Tipu* is in the same clade with different species of two genera Centrolobium and Pterocarpus (Fig 18).

**Fig 17.**
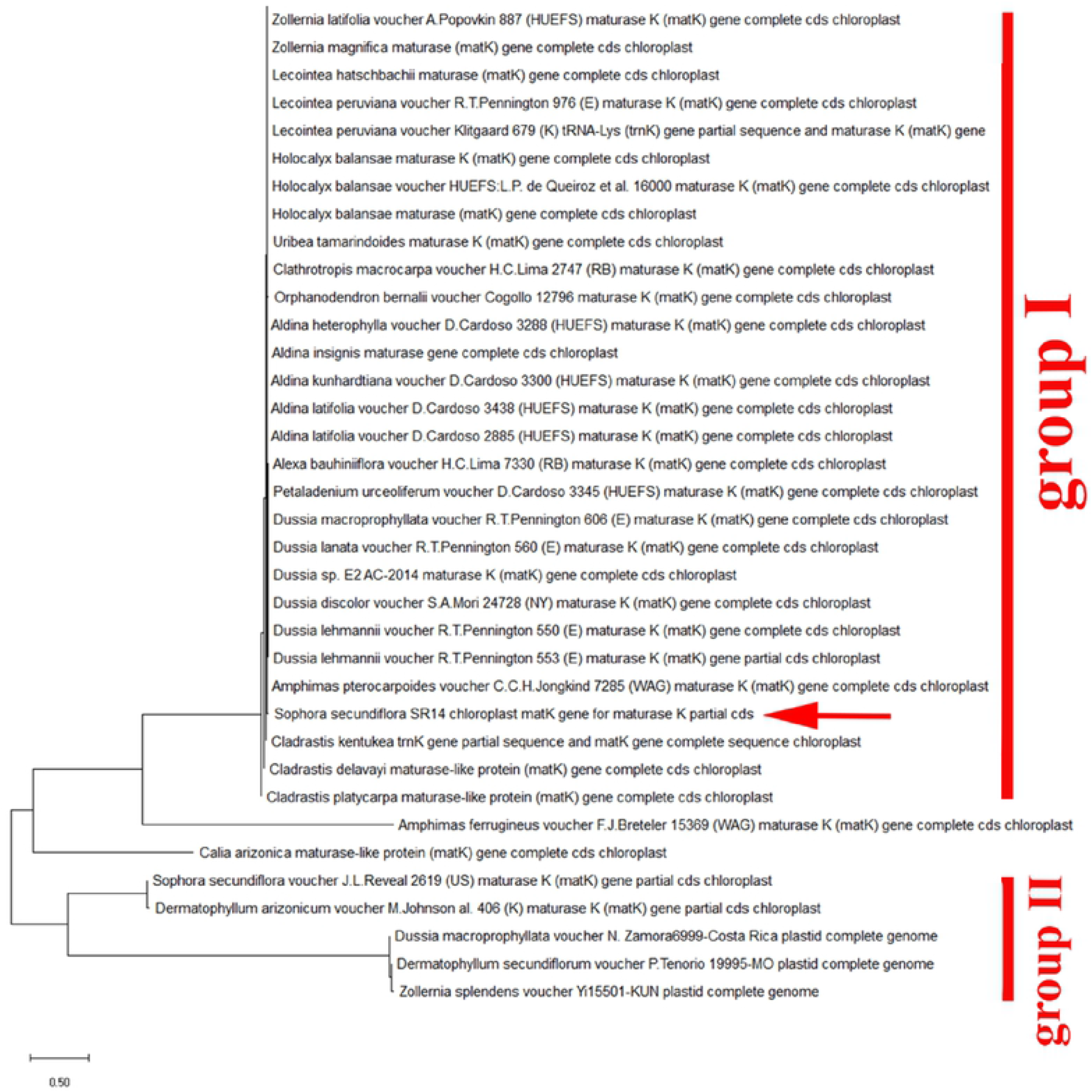
Phylogenetic tree analysis and the evolutionary distances of *Sophora secundiflora*

**Fig 18:**
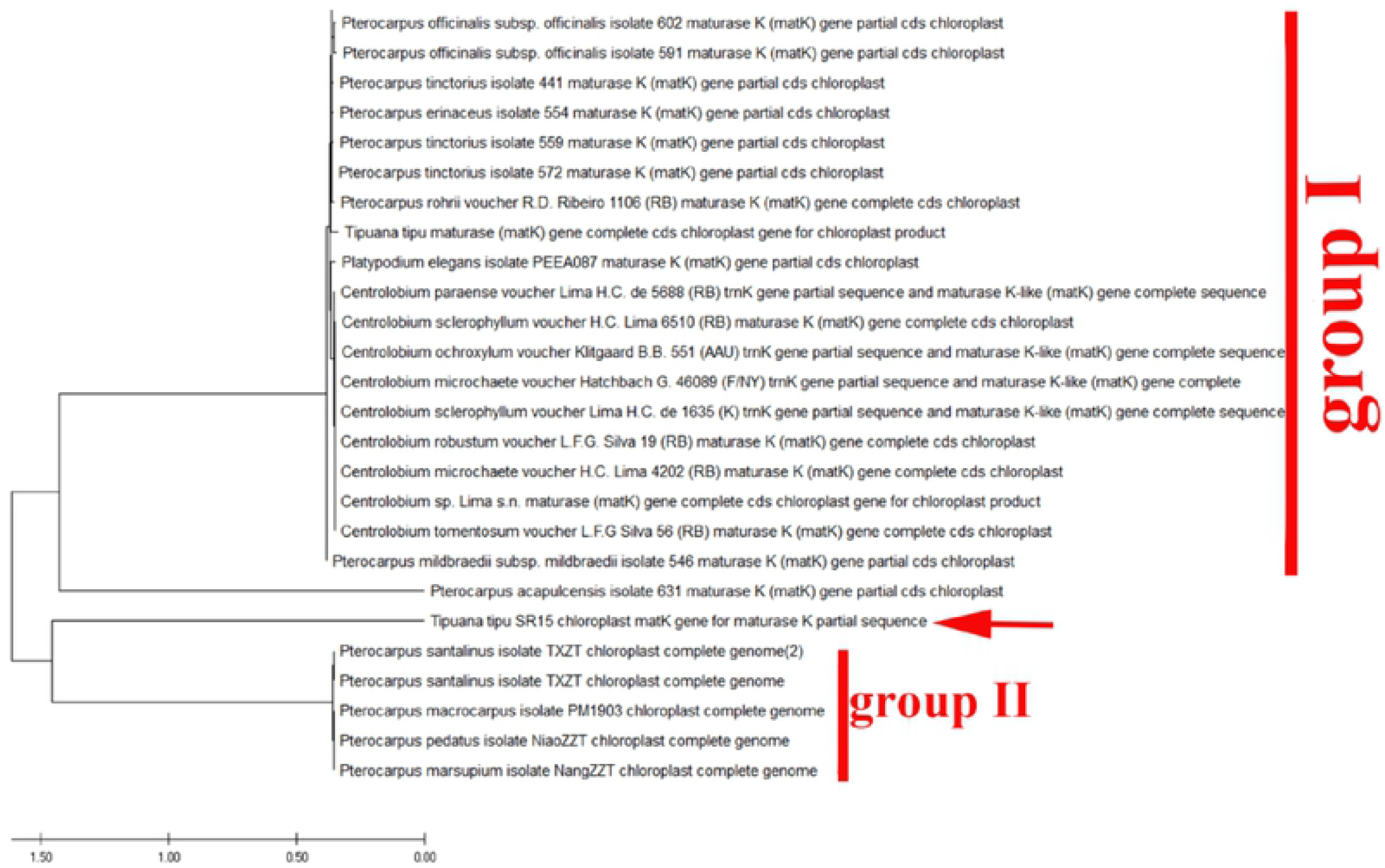
Phylogenetic tree analysis and the evolutionary distances of *Tipuana Tipu*

Additionally, highly Fabaceae species in the current research were detected to have a unique sequence in the *MatK* gene. These results will offer a valuable way to authenticate various *MatK* species. *MatK* sequence created in this analysis will be applied to construct reference sequence libraries, and the sequences extracted from samples with particular identity classifications will be utilized to search the database. Lastly, utilizing BLAST1 and the closest genetic distance approach, we will be able to define the species identities of the query sequences based on these data. In the dataset of MatK, the nearest genetic distance approach achieved 99.68% to 96.45% identification accomplishment rates at the species level for “BLAST1” and distance discrimination methodology, respectively, with no equivocal identification at the genus level. The planned barcoding portion of *MatK* is about 760 base pairs in *Fabaceae*. The phylogenetic tree (Fig 19) consists of two clades, the first clade comprising *(Enterolobium contortisiliquum, Albizia lebbek), Acacia saligna, Leucaena leucocephala, Dichrostachys Cinerea, (Delonix regia, Parkinsonia aculeata), (Senna surattensis, Cassia fistula, Cassia javanica)* and *Schotia brachypetala* were more closely to each other, respectively. The other four species of *Erythrina humeana with Sophora secundiflora and (Dalbergia Sissoo, Tipuana Tipu)* constituted the second clade.

**Fig 19:**
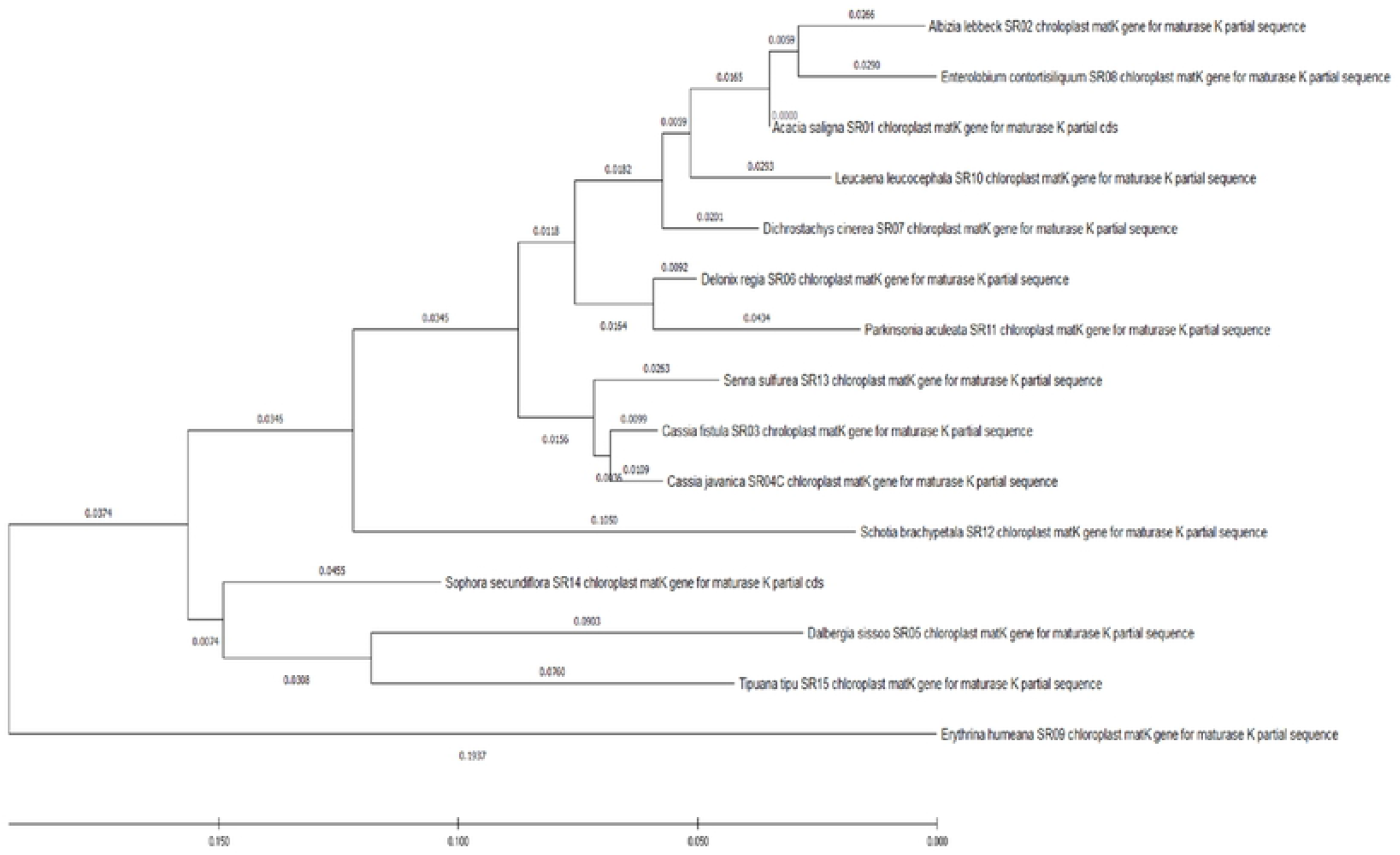
Evolutionary analysis of legume tree species grown in Egypt in this study using *Matk* gene by Maximum Likelihood method.

## 4. Discussion

Because plant genomes include several copies of MatK sequences, it’s unclear if the sequence obtained by PCR will be balanced and representative [35]. As a result, we suggest MatK as a potential barcode sequence in the Fabaceae family, as well as a wider range of plant species. Utilizing *MatK* as a DNA barcode would extend our knowledge of phylogenetics and population genetics in Fabaceae species. We also recommend that *MatK* be used as a DNA barcode sequence to overcome difficulties in Fabaceae genus and species categorization. MatK might serve as a starting point for quality control and assurance of plant materials utilized in research, manufacturing, customs, and forensics.

The *MatK* was discovered to be a necessarily variable DNA region between Fabaceae species as determined by genetic divergences, and it demonstrated a greater potential of effective discriminating. *MatK* can be a powerful taxonomic marker for identifying species and resolving taxonomic issues [36]. For instance, the *MatK* sequence of *Enterolobium contortisiliquum* is highly like *Albizia lebbek*, so our results indicate that in the genus *Cassia*, in which the species were poorly graded, *MatK* was still able to distinguish among some confusing species. The evolutionary distances for the 15 plant species that were separated into distinct clades were analyzed using the maximum likelihood and neighbor-joining methods, which discriminated most of the species better than previous techniques [37].

The identification by *MatK* region paired with morphological recognition 100% to species (Fig 19) level; for the set of plants studied, it appears to be an accurate approximation of species identification using this one locus. Short sequence, universality, and unique identifiers are three features of a common barcode [38, 39]. According to our results of sequence length and composition of *MatK* barcode gene for the 15 plant species, *MatK* regions have high rate of nucleotide substitutions as showed by [40] or the locus remodelling ring [41]. Alternate primer sequences may increase the success rate of *MatK* amplification for some of the current taxa, making it a barcoding locus. The species in which the *MatK* region is amplified, however, had wide taxonomic coverage in the Fabaceae family, indicating that the locus’ conserved sequence is notable.

Consequently, the partial amplification sequence of *MatK* was further utilized to investigate the evolutionary linkage of the selected plants. The evolutionary distances between the 15 plant species were divided into two clades using the neighbor-joining approach., the first clade comprising *(Enterolobium contortisiliquum, Albizia lebbek), Acacia saligna, Leucaena leucocephala, Dichrostachys Cinerea, (Delonix regia, Parkinsonia aculeata), (Senna surattensis, Cassia fistula, Cassia javanica)* and *Schotia brachypetala* which were more closely to each other, respectively. The remaining four species of *Erythrina humeana, (Sophora secundiflora, Dalbergia Sissoo, Tipuana Tipu)* constituted the second clade. The results are encouraging, which give a backbone of knowledge in the data set; As additional species become accessible, more research for species resolution of a genus may be undertaken.

## Author Contributions

Data curation: Aly Z. Abdelsalam, Samar M.A. Rabie, Houssam El-Din M.F. El-wakeel, Amera F. Zaitoun and Mohamed E. Hasan & Nader R. Abdelsalam; Formal analysis: Samar M.A. Rabie, Houssam El-Din M.F. El-wakeel, Amera F. Zaitoun, Hesham M. Aly and Mohamed E. Hasan & Nader R. Abdelsalam; Funding acquisition, Samar M.A. Rabie, Hesham M. Aly, Houssam El-Din M.F. El-wakeel, Amera F. Zaitoun and Mohamed E. Hasan, Alaa A. Hemeida & Nader R. Abdelsalam; Methodology, Samar M.A. Rabie, Houssam El-Din M.F. El-wakeel, Amera F. Zaitoun, Hesham M. Aly and Mohamed E. Hasan, Alaa A. Hemeida & Nader R. Abdelsalam; Resources, Samar M.A. Rabie, Houssam El-Din M.F. El-wakeel, Amera F. Zaitoun; Writing – original draft, Mohamed E. Hasan, Samar M.A. Rabie, Houssam El-Din M.F. El-wakeel, Amera F. Zaitoun, Mohamed E. Hasan, Alaa A. Hemeida and Nader R. Abdelsalam; Writing – review & editing, Alaa A. Hemeida and Nader R. Abdelsalam.

## Data Availability Statement

The data used to support the findings of this study are included within the article.

## Acknowledgment

The authors gratefully acknowledge their universities for their support

## Compliance with Ethical Standards

This study complies with relevant institutional, national, and international guidelines and legislation.

“This study does not contain any studies with human participants or animals performed by any of the authors.”

